# Well-resolved phylogeny supports repeated evolution of keel flowers as a synergistic contributor to papilionoid legume diversification

**DOI:** 10.1101/2024.10.07.617045

**Authors:** Liming Cai, Domingos Cardoso, Lydia G. Tressel, Chaehee Lee, Bikash Shrestha, In-Su Choi, Haroldo C. de Lima, Luciano P. de Queiroz, Tracey A. Ruhlman, Robert K. Jansen, Martin F. Wojciechowski

**Author notes:** Authors for correspondence: Liming Cai, Domingos Cardoso, Martin F. Wojciechowski.

## Abstract

- The butterfly-shaped keel flower is a highly successful floral form in angiosperms. These flowers steer the mechanical interaction with bees and thus are hypothesized to accelerate pollinator-driven diversification. The exceptionally labile evolution of keel flowers in Papilionoideae (Fabaceae) provides a suitable system to test this hypothesis.
- Using 1,456 low-copy nuclear loci, we confidently resolve the early divergence history of Papilionoideae. Constrained by this backbone phylogeny, we generated a time tree for 3,326 Fabales to evaluate the tempo and mode of diversification within a state-dependent evolutionary framework.
- The first keel flowers emerged around 59.0 Ma in Papilionoideae, predating the earliest fossil by 3–4 million years. The Miocene diversification of Papilionoideae coincided with rapid evolution of keel flowers. At least six independent origins and thirty-two losses of keel flowers were identified in Papilionoideae, Cercidoideae, and Polygalaceae. However, the state-dependent diversification model was not favored.
- Lack of radiation associated with keel flowers suggests that diversification within Papilionoideae was not solely driven by pollinator-mediated selection, but instead an outcome of the synergistic effects of multiple innovations including nitrogen fixation and chemical defense as well as dispersal into subtropical and temperate regions.

## INTRODUCTION

Animal pollination is one of the most striking features of angiosperms (Kay *et al*., 2006). Adaptive radiations associated with floral specialization are among the most widely discussed concepts in evolutionary biology since the era of Darwin (1862). Comparative analyses revealed several floral traits including zygomorphy that are frequently linked with species-rich clades (Sargent, 2004; O’Meara *et al*., 2016). Further ecological investigations provided mechanistic explanation through which zygomorphy can effectively guide pollinator behavior and exclude less effective pollinators via visual and structural aids (Neal *et al*., 1998; Sargent, 2004; Yoder *et al.,* 2020). Variations in the spatial organization of the perianth, stamen, stigma, and nectar spur can further localize pollen deposition spots, thus increasing reproductive isolation (Ree, 2005; Muchhala & Potts, 2007). Moreover, higher pollination efficiency and precision in zygomorphic flowers may bolster species persistence by reducing extinction risks (Kay *et al*., 2006). However, additional intrinsic (e.g., chemical defenses) and extrinsic factors (e.g., biome shifts) are often associated with lineages with specialized floral syndromes. This makes it challenging to understand the relative role of key innovations like floral specialization in promoting diversification. Indeed, about half of the comparative studies published to date did not identify a significant correlation between reproductive traits and increased diversification rate (Helmstetter *et al.,* 2023). Therefore, there is a growing trend in the field to recognize the synergistic multidriver hypothesis of angiosperm radiation rather than the key innovation hypothesis (Magallón & Castillo, 2009; Augusto *et al.,* 2014; Donoghue & Sanderson, 2015). Along these lines, papilionoid legumes (subfamily Papilionoideae, Fabaceae) are especially suitable to test key innovation versus synergistic hypotheses given their highly specialized keel flower and a suite of defense traits and biome shift history known to impact legume diversification (Lewis *et al*., 2005).

Legumes are among the most ecologically and evolutionarily successful plant families on Earth. With approximately 23,000 species and 800 genera, they have defined biomes from tropical savannas to temperate sagebrush steppes (Schrire *et al*., 2005; Lewis *et al*., 2005; LPWG, 2017; Legume Data Portal, https://www.legumedata.org/). Members of Fabaceae subfamilies often exhibit lineage-specific floral traits tailored to particular pollinators, such as the showy stamens in the Mimosoid clade for bees, the night-blooming Caesalpinioideae for moths and bats, and the distinctive butterfly-like keel flowers in Papilionoideae, which are closely linked to bee pollination (Arroyo, 1981). Keel flowers are zygomorphic with one highly differentiated standard petal, two wing petals, and a keel composed of two partially to fully fused petals enclosing the reproductive organs (Westerkamp, 1997). Connation among floral components, including stamens and keel petals, is common. They are most commonly found in two species-rich clades within Fabales—Papilionoideae and Polygalaceae, although with different ontogenetic origins (Bello *et al*., 2010). The standard serves as a visual attraction while the wings form the landing complex for pollinators (Westerkamp & Weber, 1999). The zygomorphic form and fused stamens concealed by the keel can effectively restrict the direction of entry by pollinators and aid with precise pollen placement (Neal *et al*., 1998). Furthermore, many keel flowers exhibit primary and secondary pollination mechanisms such as valvular, pump, explosive, and brush mechanisms to prevent pollen waste and deposit pollen on particular bee body parts (Leppik, 1966; Etcheverry *et al*., 2012; Uluer *et al*., 2022). Loss of keel flower traits in Papilionoideae, including actinomorphy or even exposure of reproductive organs, can lead to rapid transitions to hummingbird or bat pollination (Arroyo, 1981; Neill, 1987). Moreover, several angiosperm families independently evolved the keel flower architecture, including Ranunculaceae, Geraniaceae, Balsaminaceae, and Commelinaceae (Westerkamp, 1997). They exhibit structural and functional convergence for bee pollination, although with divergent developmental pathways (Tucker, 1999, 2003; Prenner, 2004; Prenner & Klitgaard, 2008; Bello *et al.,* 2012). Therefore, the evolution and maintenance of keel flowers is considered to be an outcome of pollinator-mediated selection with the potential to promote diversification (Grant, 1949; Stebbins, 1970; Rosas-Guerrero *et al*., 2014; Stewart *et al*., 2022).

To rigorously assess trait-driven diversification, a system with high trait variability is essential for robust comparative hypothesis testing (Ricklefs & Renner, 1994, 2000; Donoghue & Sanderson, 2015). The remarkable diversity in floral symmetry and architecture in the Papilionoideae offers an ideal model for such investigations. Although keel flowers are considered an essential feature and a hallmark of Papilionoideae (Polhill *et al*., 1981), there is considerable variation in floral symmetry in early diverging clades. For example, members from the tribes Sophoreae and Swartzieae are frequently actinomorphic, with five petals undifferentiated or even lost to various degrees (Pennington *et al*., 2000; Cardoso *et al*., 2012, 2013b). Such morphological variability is comparable to that found in the Caesalpinioideae, suggesting an exploratory phase when a broader range of ontogenetic and morphological pathways were realized (Leite *et al*., 2015). These genera are typically species-poor and more than 80% have no more than ten extant species (Plants of the World Online – POWO; https://powo.science.kew.org). In contrast, the keel flower is highly constrained among the large non-protein-amino-acid accumulating (NPAAA) clade (ca. 10,200 species), which is mostly known for economically important crops like soybeans, peas, lentils, and clovers (Choi *et al*., 2022). These general observations have prompted our speculation that the evolution of keel flowers may be associated with increased diversification rates in Papilionoideae.

Testing these macroevolutionary hypotheses requires a robust and densely sampled phylogeny. Over the past three decades, a general phylogenetic framework for Fabaceae has been developed thanks to a tremendous network of global collaboration (LPWG, 2013, 2017). Six distinct subfamilies are recognized by the Legume Phylogeny Working Group (LPWG), with Papilionoideae as the most species-rich subfamily comprising 14,000 species in 503 genera (Lewis *et al*., 2005; LPWG, 2017). However, the LPWG’s (2017) phylogeny is largely based on the plastid gene *matK*, which has obvious limitations when dealing with rapid radiations or complex evolutionary scenarios such as hybridization (Barrett *et al*., 2014; Morales-Briones *et al*., 2021). In Papilionoideae, studies based on *matK* and complete plastomes have consistently defined 22 well-supported clades but the relationships among these clades remain poorly resolved (Wojciechowski *et al*., 2004; McMahon & Sanderson, 2006; Cardoso *et al*., 2012, 2013b; Zhang *et al*., 2020; Choi *et al*., 2022). Most significantly, Swartzieae and the ADA clade have been individually proposed as sister to all remaining Papilionoideae. Given that Swartzieae contains mostly non-keel flowered taxa with frequent petal losses whereas the ADA clade contains largely keel flowered ones (Torke & Schaal, 2008; Cardoso *et al*., 2015), their placement will have a strong bearing on the timing and evolutionary trajectory of keel flowers. Furthermore, many of the 22 clades are characterized by tropical trees that are scarce in distribution and underrepresented in previous studies. These include species-poor clades such as the African *Amphimas* Pierre ex Harms, the Amazonian *Aldina* Endl., and the North American *Dermatophyllum* Scheele (Cardoso *et al*., 2013). Along these lines, recent advances in legume phylogeny using multi-locus nuclear datasets have promising applications in resolving complex evolutionary histories (Koenen *et al*., 2021; Zhao *et al*., 2021) and inspired us to apply this technique to address the recalcitrant relationships within Papilionoideae.

Here, we examined the primary versus synergistic role of keel flower evolution in Papilionoideae evolution. We first generated a phylogenomic dataset of 1,456 low-copy genes from 287 species with a focus on recalcitrant clades. Using both coalescent and concatenation methods, we inferred a robust species tree and consistently resolved the early divergence history of Papilionoideae. This nuclear phylogeny was subsequently used as the backbone to reconstruct a 3,326-species time tree of Fabales to explore the evolution of keel flowers under a broader taxonomic context in Papilionoideae, Cercidoideae, and Polygalaceae. Through ancestral character reconstruction and diversification rate inference, we evaluated the degree to which this rate variation could be attributed to keel flowers in light of a key innovation hypothesis. We considered lack of support for this hypothesis if the trait-dependent diversification model was not preferred and discussed our results in the context of the ecology and biogeography of legume species to identify the processes that account for their extreme diversity.

## MATERIALS AND METHODS

### Taxon sampling

Our taxon sampling included 279 species broadly distributed across Papilionoideae, plus eight species from other legume subfamilies as outgroups (Tables S1–2). The dataset comprised an aggregation of 47 newly generated transcriptomes and 32 low-pass genomic sequences (Table S1) as well as published sequences from 208 species from Zhao *et al*. (2021) to broaden taxon sampling (Table S2). Newly generated transcriptomes and genomic sequences were mostly from morphologically and evolutionarily representative genera within Papilionoideae (Cardoso *et al*., 2012; Choi *et al*., 2022). Collectively the dataset represented 19 of the 22 well-supported clades consistently recognized in the subfamily (Cardoso *et al*., 2012; Cardoso *et al*., 2013b; Choi *et al*., 2022). The three clades we did not include are *Amphimas*, *Aldina*, and the recently described *Cabari* (Gregório *et al*., 2024). All newly generated sequences were derived from plant materials collected in the field or grown in the University of Texas at Austin greenhouses (Table S1). Voucher specimens were deposited in the herbaria of the Rio de Janeiro Botanic Garden (RB), State University at Feira de Santana (HUEFS), and the Billie L. Turner Plant Resources Center (TEX-LL). For DNA and RNA extraction, the methods described previously by Lee *et al*. (2021) and Zhang *et al*. (2013) were employed, respectively.

### RNA sequencing and transcriptome assembly

Total RNA samples were shipped to GENEWIZ (South Plainfield, NJ, USA) for ribosomal RNA (rRNA) depletion, library preparation, and sequencing of approximately 25 million paired-end reads on an Illumina HiSeq platform (Illumina, San Diego, CA). We assessed the quality of raw 150 bp paired-end reads using FastQC v.0.11.5 (www.bioinformatics.babraham.ac.uk/projects/fastqc) and removed filtered adapters and low-quality reads using TrimGalore v.0.5.0 (https://github.com/FelixKrueger/TrimGalore). The rRNA was removed from the merged trimmed reads using SortMeRNA v.2.1 (Kopylova *et al*., 2012) using eight rRNA databases for prokaryotes, archaea, and eukaryotes. Clean RNA-Seq reads were assembled *de novo* using Trinity v.2.8.4 (Grabherr *et al*., 2011), and the assembled transcripts were subjected to the assessment of completeness and redundancy using BUSCO v.4.1.0 (Benchmarking Universal Single-Copy Orthologs; Simão *et al*., 2015) with the eudicots_odb10.2019-11-20 database. The number of raw reads and BUSCO evaluation results are provided in Table S1. All newly generated reads are available on GenBank Sequence Read Archive under Bioproject PRJNA1120003 (https://www.ncbi.nlm.nih.gov/sra/PRJNA1120003).

### Low-copy nuclear dataset assembly

We used the 1,559 low-copy nuclear loci identified by Zhao *et al*. (2021) for our phylogenetic reconstruction. To recover these loci in our transcriptomes, we used the profile hidden Markov models implemented in HMMER v3.3.2 (Johnson *et al.,* 2010) to identify orthologs. Briefly, a DNA profile database was built for each locus using the function ‘hmmbuild’ based on the DNA alignments from Zhao *et al*. (2021). These profiles were then searched against the target transcriptomes with an e-value threshold of 1e^-50^ using ‘hmmsearch’. We then extracted the best hit from each species as the candidate ortholog using a custom Python script (all scripts available on GitHub repository https://github.com/lmcai/Papilionoideae_phylogenomics).

To recover the same set of loci from our genome sequence data, we developed a new function of the PhyloHerb pipeline (Cai *et al*., 2022) to automatically extract nuclear loci from low-pass genome sequencing data. Briefly, PhyloHerb accomplishes this in three steps — mapping, assembly, and scaffolding. First, it mapped reads to the reference sequence using the very-sensitive mode of Bowtie v2.3.4.1 (Langmead & Salzberg, 2012). Mapped reads were then assembled into contigs using SPAdes v3.8.0 under the default setting (Bankevich *et al*., 2012). The contigs were scaffolded into full-length loci based on syntenic information compared to the reference using BLAST with an e-value threshold of 1e^-40^. These functions were integrated as the ‘-m assemb’ and ‘-m ortho -nuc’ functions in PhyloHerb v1.1. For details, see the GitHub page (https://github.com/lmcai/PhyloHerb/#iv-retrieve-low-copy-nuclear-genes). Finally, for quality control purposes, we also estimated the nuclear genome coverage of genome sequences using Jellyfish v2.3.0 with a kmer size of 21 (Marçais & Kingsford, 2011).

### Sequence alignment, dataset cleaning, and phylogenetic inference

Sequence alignment of each locus was conducted in MAFFT v7.245 (Katoh & Standley, 2013) under the default settings. The aligned sequences were subjected to three rounds of stringent cleaning to remove excessive missing data and potential paralogs. First, we used the ‘-missing’ function in PhyloHerb to remove sequences with >70% ambiguous sites including gaps in each locus. Second, a phylogeny-based ortholog identification method from Yang and Smith (2014) was used to remove potential paralogs. To accomplish this, a maximum likelihood (ML) phylogeny was inferred for each gene using the DNA alignments from the previous cleaning step. The best substitution model was automatically determined by ModelFinder implemented in IQ-TREE v.2.2.2.6 (Minh *et al*., 2020). Branches that were more than fifteen times the mean branch length were pruned from the resulting trees using a custom Python script ‘prune_long_branches.py’ (available on GitHub, see link above). Then, a phylogeny-based ortholog identification approach using the Python script ‘prune_paralogs_RT.py’ (Yang & Smith 2014) was applied to further remove potential contaminants of paralogs. Third, all DNA sequences were realigned using the MAFFT E-INS-i algorithm (--genafpair --maxiterate 1000) after removing spurious sequences identified in the previous steps. Sites with more than 70% missing data were removed from the alignment using the ‘-gt 0.7’ flag in trimAL v1.5.0 (Capella-Gutiérrez *et al*., 2009). We also found that the published alignments from Zhao *et al*. (2021) were site-trimmed, thus creating systematic biases for phylogenetic inference (e.g., shared gaps for sequences from Zhao *et al*. (2021) compared to species sampled by us). To accommodate this, we used a custom Python script to mask all sites in our sequences that were present in >60% of species but absent in >90% of species from Zhao *et al*. (2021). Loci with less than 199 (70%) taxa were removed from downstream phylogenomic assessments. In the end, 1,456 out of the 1,559 loci passed our screening criteria (Fig. S1).

To estimate a species tree using the multispecies-coalescent model, we first inferred individual gene trees for each of the 1,456 loci in IQ-TREE. The best substitution model was automatically determined by ModelFinder implemented in IQ-TREE and branch support was evaluated by 1000 ultrafast bootstrap replications. The species tree was inferred from 1,456 gene trees using ASTRAL-III v5.7.8 (Zhang *et al*., 2017) under the default settings. Meanwhile, to further mitigate the impact of artificially resolved gene tree branches with low support values, we used newick-utils v1.6 (Junier & Zdobnov, 2010) to collapse gene tree branches with ultrafast bootstrap support values less than 70. We then inferred a species tree with these gene trees containing polytomies in ASTRAL-III.

To infer the species tree using the concatenation methods, we first further subsampled the loci based on length and missing data. Any loci with alignments shorter than 900 base pairs (bp; 33% loci removed) or less than 227 species (80% of the total species) were excluded from the concatenation analysis, which resulted in 672 loci. The concatenated supermatrix was generated by the ‘-m conc’ function in PhyloHerb, along with a gene delineation file input into PartitionFinder 2. The best partition scheme for ML analyses employing the general time reversible (GTR) + gamma model was selected by PartitionFinder 2 using the heuristic search algorithm ‘rcluster’ (Lanfear *et al*., 2017). We then inferred an ML phylogeny of the 287 species using IQ-TREE with 100 traditional nonparametric bootstrap replications (BP) under the GTR + gamma nucleotide substitution model and linked but variable evolution rates for each partition with the ‘-p’ flag.

### Topology test

Several conflicting relationships emerged from the comparison between concatenation and coalescent analyses. To explore the origin of these conflicts and identify the best-supported phylogenetic scenarios, we conducted topology tests on six focal groups defined in Table S3. By alternating the position of the focal group and fixing the rest of the tree, we inferred an optimum phylogeny for competing scenarios in IQ-TREE. We then conducted the approximately unbiased (AU), Kishino-Hasegawa (KH), and Shimodaira-Hasegawa (SH) tests with 10,000 replicates as implemented in IQ-TREE. To compare the goodness-of-fit of alternative species trees to gene trees under the coalescent model, we conducted a pairwise likelihood ratio test (Liu *et al*., 2019). We similarly optimized the branch length of fixed topologies with alternative positions of the focal group in MP-EST v3.0 (Liu *et al*., 2010). We used the test2.sptree function from the R package Phybase v2.0 (Liu & Yu, 2010) to conduct the likelihood ratio test described in Liu *et al*. (2019). The likelihood ratio test statistic is given by *τ* = 2(*L*_Tree2_ − *L*_Tree1_), in which *L*_Tree1_ and *L*_Tree2_ are the log-likelihoods of the null and alternative hypotheses, respectively. The statistical significance (*p*-value) of the comparison is evaluated by generating a null distribution of *τ* via bootstrap sampling gene trees 100 times. Finally, to diagnose the causes of conflicting results, we examined the gene (gCF) and site concordance factors (sCF), as well as gene-wise likelihood scores implemented in IQ-TREE.

### Bayesian divergence time estimation

To infer a Bayesian time tree, we subsampled the nuclear dataset to select the most clock-like genes using SortaDate (Smith *et al*., 2018). Total tree length, root-to-tip variance, and topological congruence with the species tree were used as key indicators for phylogenetic informativeness and stable molecular rate. We selected the 30 best genes (Table S4) for dating to facilitate a reasonable running time of the Bayesian analysis, an approach comparable to a recent study of Fabaceae divergence time (Koenen *et al*., 2021). We carefully selected eleven fossils to calibrate our molecular dating analyses, including four fossils from representative eudicot lineages and seven well-recognized fossils within Fabales (Table S5). In particular, the earliest fossil fruits and leaflets of Fabaceae from the Early Paleocene (63.5 Ma) of the Denver Basin in Colorado, USA, were used to constrain the stem group Fabaceae (Lyson *et al*., 2019). Many recent fossils within Fabaceae were not included because they provide redundant constraints compared to the eleven fossils we selected. We used BEAST v.2.2.1 (Bouckaert *et al*., 2014) to estimate divergence times under the GTR substitution model, an uncorrelated relaxed molecular clock model, and a birth-death tree prior. The topology was fixed to reflect the optimum species tree inferred from the ASTRAL analysis. Fossil calibrations were set as exponential priors with the offset minimum ages aligned with the fossils described in Table S5. The root node, for which we used a normal prior at 128.6 Ma with a standard deviation of 1.0, truncated to minimum and maximum ages of 113 Ma (the Aptian–Albian boundary) and 136 Ma (the oldest known crown angiosperm fossil (Magallón *et al*., 2015). We ran analyses under the uncorrelated lognormal model for 100 million generations and confirmed convergence with Tracer v1.7.1 (Rambaut *et al*., 2018). The first 30% of the generations were discarded as burn-in before summarizing median branch lengths and substitution rates with TreeAnnotator from the BEAST package.

### Time tree inference for a 3,326-species *matK* dataset of Fabales

To evaluate the macroevolutionary trends of floral diversification in Papilionoideae under a broader taxonomic context, we generated a plastid *matK* gene dataset that included 3,106 species from 794 genera in Fabaceae (92% genus-level completeness), plus 220 additional species from all other families in Fabales and outgroups. Sequences from Choi *et al*. (2022), Legume Phylogeny Working Group (LPWG 2017), and Gregório *et al*. (2024) were consolidated to generate the most comprehensive *matK* database for Fabaceae, and combined with *matK* sequences from Polygalaceae, Surianaceae, and Quillajaceae available on GenBank, which served as outgroups in Fabales (Table S6).

As a first step of data curation, we updated the nomenclature to reflect the most recent effort from POWO, Flora e Funga do Brasil (https://floradobrasil.jbrj.gov.br/consulta; BFG 2022), and taxonomic studies from individual clades (e.g., Hughes *et al*., 2022). Second, we employed multiple rounds of phylogeny-guided cleaning for the *matK* sequence alignment. Initial alignment was conducted with the translation align option of MAFFT v.7.490 implemented in Geneious Prime 2023.1.2 (https://www.geneious.com/) and was manually inspected thereafter. For outgroup species, a separate alignment was generated using the L-INS-i algorithm in MAFFT to accommodate higher sequence divergence across families. These two alignments were then merged using the profile-profile alignment algorithm implemented in ClustalX (Thompson *et al*., 2003) and the final DNA alignment contained 3,315 aligned sites. A draft ML phylogeny was inferred under the GTR+I+G model with 1000 ultrafast bootstrap replications in IQ-TREE and we filtered the *matK* matrix accordingly to remove misidentified and rogue taxa.

To infer a robust *matK* phylogeny for divergence time estimation, we used our well-supported nuclear phylogeny to constrain the topology and node ages of major clades in the *matK* dataset. This is necessary because the slow substitution rate of *matK* may result in contentious relationships and the lack of sequence divergence among close relatives may impact ancestral state estimation and divergence time estimation. As a result, relationships *between* the following clades were constrained: the ADA, Swartzieae, Cladrastis, Vataireoid, Exostyleae, Dalbergioid s.l., Genistoid s.l., Andira, Baphieae, and NPAAA. Relationships *within* each clade and *between* all subfamilies and families remained as-is. With this partially fixed topology, the ML phylogeny was inferred under the best fitting model in IQ-TREE.

Secondary calibration points of thirty nodes from the Bayesian time tree (Table S7) were used to infer the *matK* gene time tree under the penalized likelihood method implemented in TreePL v1.0 (Smith & O’Meara, 2012). These calibration points were imposed as both maximum and minimum age constraints to fix node ages. We then primed and cross-validated our data to establish the best smoothing parameter (smooth = 10) for our dataset. Finally, we conducted a thorough search to infer the best time tree.

### Testing state-dependent diversification analysis

To explore the diversification dynamics across Fabales, we performed a Bayesian analysis of macroevolutionary mixtures using the program BAMM v2.5.0 (Rabosky, 2014), which assumed clade- and time-dependent diversification scenarios to detect shifts in diversification rates. Reversible-jump Markov chain Monte Carlo was run for 20 million generations on the complete Fabales phylogeny and sampled every 1,000 generations. Posterior probabilities of rate shifts were summarized with the R package BAMMtools v.2.1.6 (Rabosky *et al*., 2014) and visualized using a customized script from Padfield *et al*. (2024).

To explore whether the evolution of keel flowers drives pollinator-associated diversification in clades like Papilionoideae (Fabaceae) and Polygaleae (Polygalaceae), we estimated the divergence time history of Fabales under the state-dependent speciation and extinction (SSE) models. First, we coded the floral type of all species as either keel or non-keel flowers defined as above by Westerkamp (1997; Table S6). To infer the sampling ratio, we used the powoGenera function from the R package expowo (Zuanny *et al*., 2024) to obtain the number of species for each genus based on the Plants of the World Online database (https://powo.science.kew.org; Table S8). The total species number for the keel flower clades and non-keel flower clades was enumerated to calculate the sampling ratio for these two groups. Finally, we inferred the diversification rate under the state-dependent and state-independent models using the R package hisse v2.1.8 (Beaulieu & O’Meara, 2016). We estimated rate parameters for four types of models (Table S9): (1) the Binary State Speciation and Extinction model (BiSSE), which allows state-dependent diversification (i.e., keel versus non-keel) but does not include hidden states; (2) the Hidden State Speciation and Extinction model (HiSSE), which allow both state-dependent diversification as well as hidden states; (3) CID-2 and CID-4 models, which are state-independent but allow differential diversification associated with the hidden states to serve as the null model for BiSSE and HiSSE, respectively; and (4) the trait-free, hidden state-only rate model MiSSE for tip rate estimation (Maddison *et al*., 2007; Beaulieu & O’Meara, 2016; Vasconcelos *et al*., 2022). For the HiSSE analyses, we included models with the number of hidden states from two to four, which encompassed all combinations of hidden states supported in the hisse package. To reconstruct the ancestral states of keel flowers, we used the function ‘hisse’ followed by ‘MarginReconHiSSE’ to infer the speciation and extinction rates as well as ancestral states under the standard HiSSE model with two hidden states. For the MiSSE analyses, the optimum number of hidden states was tested for all possible configurations ranging from 1 to 26. The best model with the lowest AIC was selected for downstream analysis. We then applied the ‘MarginReconMiSSE’ function to estimate the tip rates and evaluate the correlation between floral architecture and diversification. We conducted a phylogenetic ANOVA test using the phylANOVA function from the R package phytools (Revell, 2012) to compare tip rates between keel and non-keel like species. Besides testing SSE model fitness using the entire Fabales phylogeny, we also applied the BiSSE, HiSSE, CID-2, and CID-4 models to a subsampled phylogeny that only included Papilionoideae species. Together, this resulted in a total of 11 different analyses (Table S9).

## RESULTS

### An efficient phylogenetic data assembly pipeline based on a curated set of loci

BUSCO assessments for the assembled transcriptomes showed a high level of completeness, with the percentage of complete orthologous genes ranging from 70.3% to 91.1% (Table S1). The final multi-locus phylogenetic dataset included 1,456 genes with a mean length of 1,003 bp after trimming (Data S1). These loci on average contained 253 (85%) species and the concatenated DNA dataset consisted of 941,354 sites with a gene and character occupancy of 91.5% and 83.0%, respectively. None of our newly sequenced species exhibited exceptional levels of missing data (<30% missing; Table S1), but three species added from Zhao *et al*. (2021), which included *Pisum sativum* L., *Hesperothamnus pentaphyllus* (Harms) Harms, and *Euchresta japonica* Hook. f. ex Regel, contained the highest level of missing data (69–85% missing; Table S2).

Our comprehensive, genus-level taxon sampling represents multiple data collection strategies, including high-quality reference genomes, RNA-seq, and low-pass genome sequencing (i.e., genome skimming). Among these three approaches, our *in-silico* extraction of phylogenetic markers from genomic sequencing data using PhyloHerb represents a cost-effective approach to construct a multi-locus phylogeny, eliminating the need for target enrichment library preparation. Genomic sequencing of the 32 Papilionoideae taxa yielded 25–30 million 150 bp paired-end reads and 1–15× nuclear coverage according to k-mer distribution (Table S10; Data S2). In the final DNA matrix, the level of missing data in these 32 species was not statistically different from transcriptomic data (Welch two sample t-test *p*-value = 0.08). Even for species with less than 2× nuclear genome coverage, at least 1,075 (74%) of the targeted loci were recovered (Table S10). Computationally, PhyloHerb is also highly accessible. Extracting ∼1,500 loci from each species on average requires 903.2s CPU time and 2.9 GB peak memory including assembly and ortholog identification, which can be completed on most laptops.

### Multi-locus nuclear dataset resolves contentious relationships in Papilionoideae

To accommodate for incomplete lineage sorting, we used both coalescent and concatenation methods for phylogenetic inference. The coalescent-based phylogeny estimated by ASTRAL was largely congruent with the concatenation-based phylogeny inferred by IQ-TREE and 92.6% and 89.1% of the nodes received maximum support (100 BP; 1.0 posterior probability, PP), respectively (Fig. 1).

**Figure 1.**
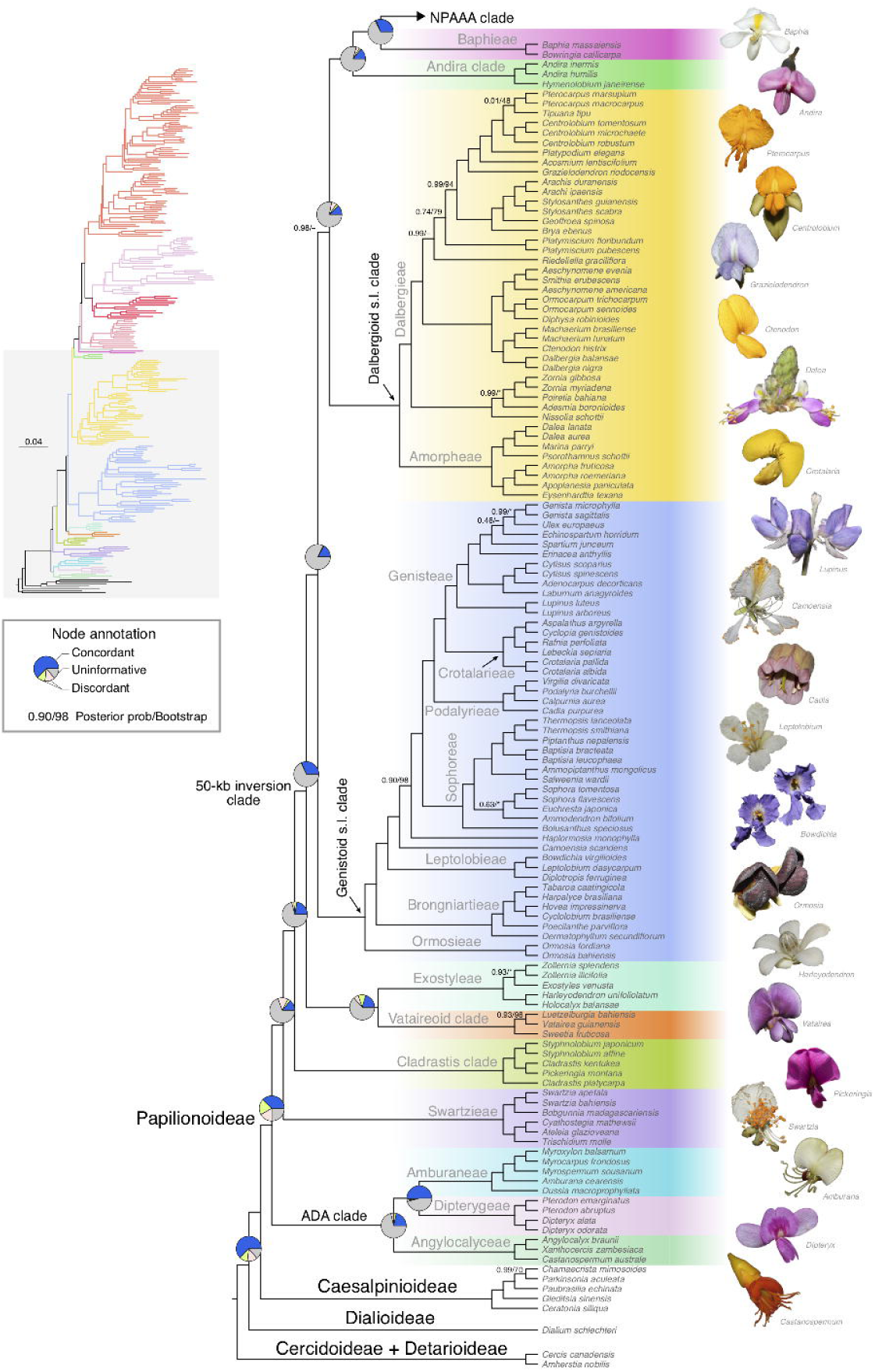
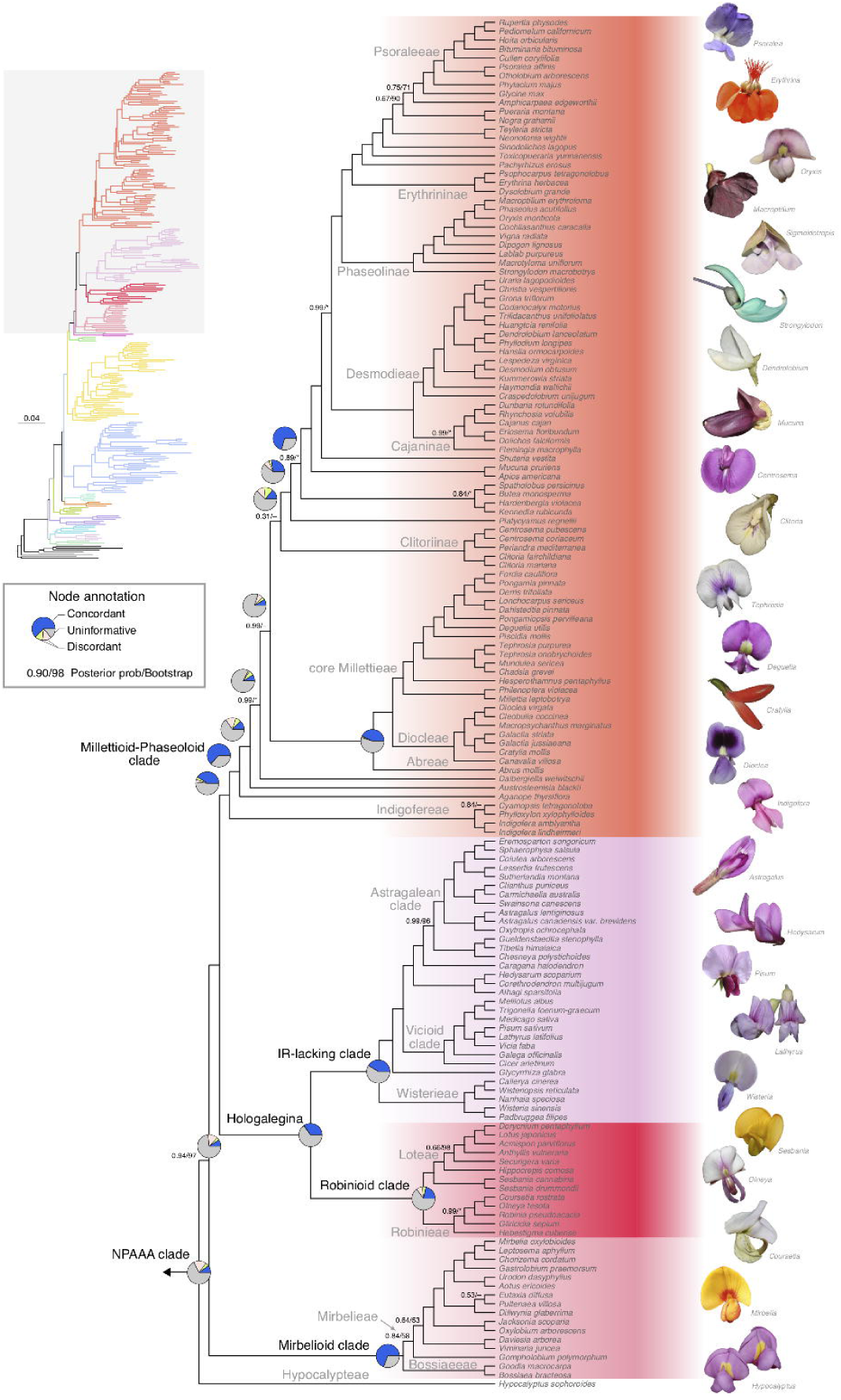
Phylogeny of Papilionoideae based on low-copy nuclear gene sequences and inferred from the multispecies coalescent model implemented in ASTRAL-III. Branch supports indicate posterior probability from ASTRAL and bootstrap replicates support from IQ-TREE, respectively. Only branches that are not maximally supported are labeled with conflicting topologies marked by ‘–’. Pie charts on each node represent gene concordance factors for the proportion of concordant (blue), discordant (green and pink), and noninformative (grey) genes. The phylogram in the left margin illustrates branch length in mutational units. The photos (by Domingos Cardoso, except *Pickeringia* by Martin Wojciechowski, *Grazielodendron* by Claudio Nicoletti, *Baphia* by Warren McCleland (iNaturalist, https://www.inaturalist.org/photos/108404418), *Hypocalyptus* by Brian du Preez (iNaturalist, https://www.inaturalist.org/photos/60072646), *Mirbelia* by Nina Kerr (iNaturalist, https://www.inaturalist.org/photos/345474574), *Pisum* by Michela Papa (iNaturalist, https://inaturalist.ca/photos/372960709), and *Psoralea* by katrivier (iNaturalist, https://inaturalist.ca/photos/262747553) illustrate the remarkable diversity of floral architecture within the Papilionoideae.

Within Papilionoideae, the monophyly of 19 well-recognized clades was fully supported by both coalescent and concatenation analyses (Fig. 1; Fig. S2). Relationships among these clades were confidently resolved including those of recalcitrant lineages such as *Dermatophyllum, Sesbania,* and the Andira clade (Fig. 1). The ADA, Swartzieae and Cladrastis clades formed successive sister groups to all remaining members of Papilionoideae — the 50-kb inversion clade (Fig. 1). Within the 50-kb inversion clade, the Vataireoid and Exostyleae clades together formed a well-supported clade (100 BP; 1.0 PP) sister to all other groups. The monophyly of the Vataireoid and Exostyleae clades was strongly favored by both the AU test using the concatenated matrix and the likelihood ratio test based on the multispecies coalescent (MSC) model. The alternative topology suggested by Zhao *et al*. (2021), where these two groups were not monophyletic, was rejected (AU test *p-*value < 0.01; MSC likelihood ratio test *p-*value <0.01; Table S3).

One major conflict in our results is the relative placement of the Dalbergioid s.l. (1,580 spp.) and Genistoid s.l. (2,100 spp.) clades within the 50-kb inversion clade. A monophyletic Dalbergioid s.l. + Genistoid s.l. clade was recovered but poorly supported by the concatenation method (41 BP). On the other hand, the coalescent method supported the monophyly of Dalbergioid s.l. plus the Andira, Baphieae, and NPAAA clades with high confidence (0.98 PP; Fig. S2). The AU test on these two alternative topologies did not provide definitive conclusions (*p-*value >0.05; Table S3) and the gene-wise likelihood supporting each topology was approximately the same (Fig. S3). The MSC likelihood ratio test, however, rejected the monophyletic Dalbergioid s.l. + Genistoid s.l. clade (*p-*value = 0.03). Within the Genistoid s.l. clade, tribes Ormosieae, Brongniartieae, and Leptolobieae form successive sister groups to all remaining species (100 BP; 1.0 PP). *Dermatophyllum* was confidently placed as sister to Brongniartieae (88 BP; 1.0 PP). The African monospecific genus *Haplormosia* Harms was nested within the genistoids (98 BP; 0.90 PP). Within the Dalbergioid s.l. clade, all previously identified major subclades were maximally supported. Tribe Amorpheae appeared as sister to all remaining dalbergioids, and the Adesmia, Dalbergia, and Pterocarpus subclades were phylogenetically successive sisters (100 BP; >=0.99 PP).

The Andira, Baphieae, and NPAAA clades were each monophyletic and together formed a grade comprising the remaining Papilionoideae. Within the NPAAA clade, the genus *Hypocalyptus* Thunb. was sister to all other members of this clade (97 BP; 0.94 PP), while the Mirbelioid, Hologalegina, and Indigofereae + Millettioid-Phaseoloid clades formed successive sisters to Hypocalypteae. Within mirbelioids, both coalescent and concatenation methods placed lineages from tribe Bossiaeeae as sister to a poorly supported clade composed mostly of tribe Mirbelieae (<58 BP; <0.83 PP). Such low support could be attributed to the two short internal branches (1.3–1.7e^-3^ in mutation units; 2.0–3.9e^-2^ in coalescent units) separating Bossiaeeae, *Gompholobium* Sm, and the *Daviesia* Sm. + *Viminaria* Sm. clade within Mirbelieae. These rapid radiations have rendered little phylogenetic information to reconstruct this section of the phylogeny — the gene concordant factor was 16.3% for the branch separating Bossiaeeae and Mirbelieae, while 14.0% of the genes supported the other two alternative quartet topologies at this node; similarly, 16.0% of the genes supported the paraphyly of *Daviesia* + *Viminaria* and *Gompholobium*, but one of the alternative topologies was supported by 17.4% of the genes (Fig. 1; Data S3). Results from the site concordance factor largely reflected a similar trend where each of the three quartet topologies was supported by 33% of the sites (Data S3).

The Millettioid-Phaseoloid clade within the larger NPAAA clade harbored more uncertainty. The three genera *Aganope* Miq., *Austrosteenisia* R. Geesink and *Dalbergiella* Baker f. formed a well-supported basal grade within the millettioids (100 BP; 1.0 PP). The concatenation method confidently placed (100 BP) subtribe Clitoriinae — including *Centrosema* (DC.) Benth., *Periandra* Mart. ex Benth. and *Clitoria* L. — as sister to *Dalbergiella*. In contrast, the coalescent method placed Clitoriinae with a clade consisting of members from tribes Desmodieae, and Psoraleeae, and the polyphyletic Phaseoleae with low confidence (0.31 PP). A clade comprising tribes Abreae, Diocleae, Millettieae formed the next diverging group in the millettioids. The remaining millettioids included tribes Desmodieae, Psoraleeae, and the polyphyletic Phaseoleae. In both coalescent and concatenation analyses, *Platycyamus regnellii* Benth. and the *Butea* clade (*Spatholobus* Hassk., *Butea* Roxb. ex Willd., *Hardenbergia* Benth., and *Kennedia* Vent.) formed successive sister groups to all other members within this clade (100 BP; 1.0 PP). Conflicts exist in terms of the placement of the *Apios* clade (*Mucuna* Adans. and *Apios* Fabr.) and genus *Shuteria* Wight & Arn, which were monophyletic (99 BP) in the concatenation analysis but paraphyletic (0.89 PP) in the coalescent analysis.

### Evolution of keel flowers in Fabales

The robust nuclear phylogeny was used to constrain the uncertain relationships among major clades in the *matK-*based phylogeny. This broadly sampled phylogeny of Fabales included 3,326 species — 3,106 Fabaceae (15.9% species diversity), 1 Quillajaceae (100%), 7 Surianaceae (55.6%), 206 Polygalaceae (15.4%), and 6 additional eudicots for dating purposes. With fossil constraints, the time tree inferred by BEAST supported a Late Cretaceous origin of crown group Fabales at 78.0 Ma (95% HPD 71.9–88.3 Ma; Fig. S4), and an early Paleocene origin of crown group Fabaceae and Polygalaceae at 66.0 Ma (95% HPD 64.7–66.8 Ma) and 56.1 Ma (95% HPD 55.8–56.6 Ma), respectively. Within Fabaceae, we inferred an explosive diversification leading to all six subfamilies within 7 million years after the origin of the family. Crown group Papilionoideae was inferred to originate at 60.9 Ma (95% HPD 59.7–62.1 Ma; Fig. S4).

Keel flowers, which are mostly known in Papilionoideae and tribe Polygaleae (Polygalaceae), have been independently lost and gained multiple times. Ancestral character reconstruction suggested at least six independent origins of keel flowers in tribe Polygaleae (Polygalaceae), *Cercis* L. (Cercidoideae), tribe Dipterygeae (ADA clade, Papilionoideae), the MRCA of *Petaladenium* Ducke and *Dussia* Krug & Urb. ex Taub. (ADA clade, Papilionoideae), *Myrospermum* Jacq. (ADA clade, Papilionoideae), and the MRCA of the Cladrastis clade and the 50-kb inversion clade (Papilionoideae). In addition, there were two independent losses of keel flowers in Polygalaceae and thirty in Papilionoideae (summarized in Table S11; Fig. S5). These numbers, especially the losses, were strongly dependent on the phylogeny. We inferred the ancestral floral architecture of Papilionoideae to be non-keel-like, with the first keel flowers emerging in the common ancestor of the Cladrastis and the 50-kb inversion clades around 59.0 Ma (95% HPD 57.9–60.2 Ma). This is more recent than the origination of keel flowers in crown group Polygalaceae, which included *Xanthophyllum* at 62.8 Ma (95% HPD 59.6–64.9 Ma), though the exact order cannot be established with confidence due to overlapping confidence intervals.

### State-dependent diversification

The BAMM analysis inferred 18 rate shifts in Fabales and 11 of these shifts are nested within Papilionoideae (Fig. 2a, c). The diversification rate increased gradually over time in Fabales and nearly all rate shifts represented a significant acceleration in diversification rates rather than a decrease (gray lines in Fig. 2b). With few exceptions, 9 out of the 11 rate shifts in Papilionoideae took place during the Miocene. These Miocene rapid radiations involved well known examples from the mimosoids, the genera *Lupinus* L. and *Astragalus* L., Psoraleeae, and Phaseolinae (clades A, H, K, L in Fig. 2a).

**Figure 2.**
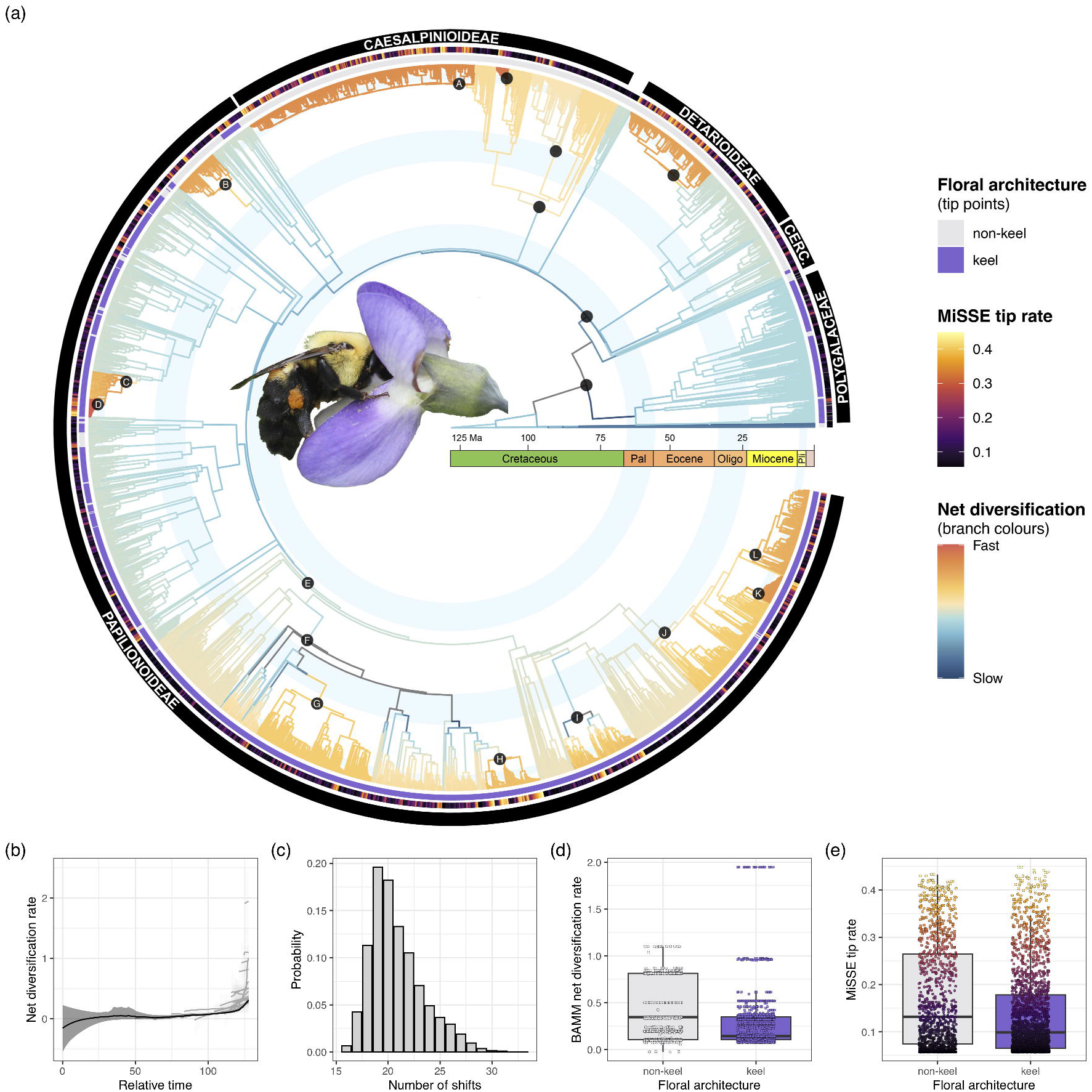
Rate heterogeneity in the diversification of Fabales. (a) Time-calibrated tree of Fabales species based on plastid *matK* gene sequences. Branch colors show net diversification rates inferred from BAMM. Circles at nodes refer to clades with significant rate shifts, which approximately align with the mimosoids (A), *Swartzia*+*Bocoa*+*Candolleodendron* (B), Genisteae (C), *Lupinus* (D), NPAAA clade (E), IR-lacking clade (F), Vicioid clade (G), *Astragalus* (H), Diocleae (I), Cajaninae+Desmodieae+Phaseolinae+Erythrininae+Psoraleeae in addition to several other smaller lineages (J), Psoraleeae (K), and Phaseolinae (L). Purple lines in the inner most ring indicate the presence of keel flowers. The middle ring represents net diversification rates at the tip estimated from the MiSSE model with 21 hidden states. Warmer colors are suggestive of higher rates. Fabaceae taxonomic groups including Cercidoideae, Detarioideae, Caesalpinioideae, and Papilionoideae are identified in the outer ring. (b) Rate-through-time plot showing increased net diversification over time. The black line represents the average across the whole tree; the gray lines represent the rate through time on parts of the tree where significant rate shifts were identified. Shaded regions represent 95% confidence intervals. (c) Posterior distribution of the number of rate shifts inferred using BAMM. (d) Variation in tip-specific diversification rates for keel-flowered and non-keel-flowered lineages based on BAMM estimate. (e) MiSSE-based tip diversification rates for keel-flowered and non-keel-flowered lineages. All rates are relative and only allow us to look at relative differences between biome preferences and parts of the phylogeny. The photo by Janet Davis shows a *Bombus griseocollis* (De Geer, 1773) bee on a *Baptisia australis* (L.) R.Br. flower.

Although most rate shifts took place in the keel-flowered Papilionoideae, our diversification rate analysis did not support the state-dependent models (Table S11). When applying the state-dependent (BiSSE and HiSSE), the state-independent (CID-2 and CID-4), and the completely trait-free models (MiSSE) to Fabales, MiSSE with 21 hidden states was significantly better than the second-best CID-4 model, with a ΔAIC of 474.75. The phylogenetic ANOVA analysis also confirmed no significant difference in the MiSSE tip rate between keel and non-keel species (*p*-value = 0.71). This conclusion held when using a subsampled phylogeny that included only Papilionoideae species. For these subsampled analyses, only BiSSE, standard HiSSE with two hidden states, CID-2, and CID-4 were tested. Among them, the state-independent CID-4 was significantly better than the second best HiSSE model with a ΔAIC of 74.67. However, diversification parameters from both BiSSE and HiSSE models suggested an orders-of-magnitude higher net diversification rate in keel-flowered clades than non-keel-flowered clades. Specifically, HiSSE suggested the net diversification rate to be 0.13 in keel-flowered clades and 5.3e^-4^ in non-keel-flowered clades; BiSSE suggested it to be 0.082 in keel-flowered clades and 7.7e^-4^ in non-keel-flowered clades.

## DISCUSSION

### Insights into the early divergence relationships within papilionoids

Previous phylogenetic studies focusing on the early divergences of papilionoid legumes have largely utilized plastid markers (Wojciechowski *et al*., 2004; Lavin *et al*., 2005; McMahon & Sanderson, 2006; Cardoso *et al*., 2012, 2013b; Zhang *et al*., 2020; Choi *et al*., 2022). These studies have consistently recognized well-supported clades but struggled in resolving inter-clade relationships. Based on 1,456 low-copy nuclear genes, we show that the ADA, Swartzieae, and Cladrastis clades are successive sister groups to the 50-kb inversion clade (Fig. 1). This result is congruent with the recent nuclear Fabaceae phylogeny from Zhao *et al*. (2021) and a multi-locus phylogenetic study of the nitrogen-fixing clade (Kates *et al*., 2024) but differs from most plastid-based phylogenies where the Swartzieae clade was placed as sister to all other Papilionoideae (Cardoso *et al*., 2012; Zhang *et al*., 2020; Choi *et al*., 2022). Our results also differ from the most recent angiosperm phylogeny based on 353 nuclear genes where the ADA and Swartzieae clades formed a monophyletic lineage sister to all other Papilionoideae (Zuntini *et al*., 2024).

The 50-kb inversion clade constitutes 95% of the remaining species diversity in Papilionoideae. Our well-resolved nuclear phylogeny provides several important insights into relationships within this clade that were inconclusive in previous plastid sequence-based studies. First, the Exostyleae and Vataireoid clades form a monophyletic group sister to the remaining members of the 50-kb inversion clade (100 BP; 1.0 PP). Their close relationship is morphologically supported by the denticulate, serrate, or occasionally spinescent leaf margin, which is highly unusual outside the NPAAA clade in Papilionoideae (Mansano *et al*., 2004; Cardoso *et al*., 2012, 2013a,b). Second, although conflicting topologies are obtained when placing the Genistoid s.l. clade, statistically significant support is only achieved for the coalescent analysis (Table S3). A monophyletic group comprising the Genistoid s.l. and Dalbergioid s.l. clades was reported by Kates *et al*. (2024) and Zuntini *et al*. (2024) but strongly rejected by our coalescent-based likelihood ratio test. Instead, our ASTRAL phylogeny placed the Genistoid s.l. clade sister to a clade comprised of the Dalbergioid s.l., Andira, Baphieae, and NPAAA clades (PP = 0.98). Within the latter clade, the Dalbergioid s.l., Andira, and Baphieae clades form successive sister groups to the large NPAAA clade — a relationship that has not been observed in any plastid sequence-based study (Cardoso *et al*., 2012, 2013b; Choi *et al*., 2022).

Third, the genus *Dermatophyllum,* comprised of four North American species, has been a clade recalcitrant to phylogenetic resolution in plastid-based investigations and often formed a polytomy with the Dalbergioid s.l., Andira, and Genistoid s.l. clades (Wojciechowski *et al*., 2004; Cardoso *et al*., 2012; Cardoso *et al*., 2013b; Kates *et al*., 2024). We confidently placed *Dermatophyllum* as sister to tribe Brongniartieae within the Genistoid s.l. clade (88 BP; 1.0 PP). This placement is supported by their shared niche preference and the ability to synthesize a class of highly distinct alkaloids: *Dermatophyllum* prefers warm temperate woodlands, chaparral, and deserts of southern United States to central Mexico, which overlaps with many Mexican Brongniartieae especially in the genus *Brongniartia* Kunth (Pennington *et al*., 2005; Ross & Crisp, 2005). The presence of quinolizidine alkaloids in *Dermatophyllum* further supports its placement within the Genistoid s.l. clade (Cardoso *et al*., 2013b; Choi *et al*., 2022). This class of alkaloids is absent in all other legumes and only sparsely distributed in eight other vascular plant families (Bisby *et al*., 1994; Kite & Pennington, 2003; Lee *et al*., 2013; Wink, 2013; Cely-Veloza *et al*., 2023). We thus demonstrate a single origin of alkaloid biosynthesis in papilionoids and a chemical synapomorphy for the Genistoid s.l. clade.

Fourth, the close relationship between the African genus *Haplormosia* and the American-Australian Brongniartieae clade proposed by Cardoso *et al*. (2017) is challenged by our new result. Although several reproductive and vegetative features are shared between these two lineages — including unifoliolate leaves; single-seeded, tardily dehiscent fruits with thick, woody valves; and stamens with distinctly dimorphic anthers (Cardoso *et al*., 2017) — the distinct biogeographic distribution makes the proposed relationship contentious. Our results instead place *Haplormosia* within a group of largely Old-World lineages, including *Camoensia* and the tribes Crotalarieae, Genisteae, Podalyrieae, and Sophoreae (98 BP; 0.90 PP). Morphologically, species from Crotalarieae, Podalyrieae, and Genistieae also exhibit unifoliolate leaves and dimorphic anthers that are alternately basifixed and dorsifixed (Schutte-Vlok & van Wyk, 2011; Le Roux & Van Wyk, 2012; du Preez *et al*., 2021).

Finally, the NPAAA clade contains a whopping ca. 10,200 species that account for nearly 70% of species diversity in papilionoids. It is sister to the predominantly African Baphieae clade with ca. 60 species (100 BP; 1.0 PP). The NPAAA clade comprises two large, strongly supported subclades, the *Hologalegina* (Robinioid+IR-lacking clade) and Indigofereae + Millettioid-Phaseoloid clades, in agreement with all previous molecular phylogenetic studies (Wojciechowski *et al*., 2004; Cardoso *et al*., 2013b; LPWG, 2017; Zhao *et al*., 2021). Within Hologalegina (ca. 5,300 spp.), the relationship between *Sesbania* and tribes Loteae and Robinieae was highly unstable with all combinations of relationships recovered using plastid data (Wojciechowski *et al*., 2004; Farruggia *et al*., 2018; Zhang *et al*., 2020; Choi *et al*., 2022). Our results confirm *Sesbania* as sister to the warm temperate, primarily northern hemisphere Loteae (100 BP; 1.0 PP), and together they form a sister group to Robinieae (100 BP; 1.0 PP). The paraphyletic relationship among *Sesbania,* Loteae, and Robinieae proposed by Zhao *et al*. (2021) is strongly rejected by our AU test (*p-*value < 0.01) and MSC likelihood ratio test (*p–*value < 0.01). Relationships within the IR-lacking clade (IRLC) align with recent studies using plastid and nuclear sequences (Choi *et al*., 2022; Zhao *et al*., 2021). Relationships within the primarily subtropical to pantropical Indigofereae + Millettioid-Phaseoloid clade (ca. 4,050 spp.) are consistent with earlier findings (Hu *et al*., 2000; Wojciechowski *et al*., 2004; Stefanović *et al*., 2009; LPWG, 2017; Oyebanji *et al*., 2020; Zhao *et al*., 2021; Choi *et al*., 2022).

### Loss and gain of keel flowers during the diversification of the papilionoids

A suite of reproductive, nutritional, and defense traits has contributed to the assemblage of present legume biodiversity (Lewis *et al*., 2005). Here we focused on the floral form in papilionoid legumes and more broadly in Fabales, which has been widely invoked to explain the radiation of many angiosperms (Sargent, 2004; Kay & Sargent, 2009; Givnish *et al*., 2015). Bilaterally symmetrical flowers such as keel flowers are strongly associated with bee pollination because the banner and wing petals can attract and orient the pollinators while the keel petals restrict the entry to pollen (Neal *et al*., 1998). The relative position of the keel, pistil, and stamens also facilitates the precise placement of pollen to pollinators and thus discourages gene flow between incipient species (Neal *et al*., 1998). Despite being a stereotyped search image for legumes, there is great lability in floral form and symmetry during the early evolution of papilionoids (Fig. 2). This provides an ideal comparative framework to test whether the evolution of keel flowers drives diversification (Ricklefs & Renner, 1994, 2000). Extending our hypothesis testing to Fabales not only broadens the investigation to another important keel lineage in Polygalaceae, but also offers evolutionary context for this highly specialized floral form.

Our well-resolved nuclear gene and densely sampled plastid *matK* gene phylogenies provide the essential evolutionary framework to address this question. Crown group Fabaceae was estimated to originate at the K-Pg boundary at 66.0 Ma (95% HPD 64.7–66.8 Ma), which is in line with previous molecular-based estimates and fossil records (Lavin *et al*., 2005; Bruneau *et al*., 2008; Cannon *et al*., 2015; Koenen *et al*., 2021; Zhao *et al*., 2021). The ancestral floral architecture of Papilionoideae is estimated to be non-keel-like, with the first keel flowers in Fabaceae emerging in the common ancestor of the Cladrastis and the 50-kb inversion clades at 59.0 Ma (95% HPD 57.9–60.2 Ma). This estimate predates the earliest known papilionoid-like flower fossil *Barnebyanthus buchananensis* Crepet & Herendeen by 3 million years (Crepet & Herendeen, 1992) and closely succeeded the evolution of keel flowers in crown group Polygalaceae at 62.8 Ma (95% HPD 59.6–64.9 Ma). However, these estimates contradict the results of a recent study which concluded that papilionoid keel flowers evolved 22.03 to 10.32 million years before they occurred in crown group Polygalaceae (Uluer *et al*., 2022).

Such a discrepancy comes mainly from the ancestral state reconstruction analysis of keel flowers themselves, but also from minor differences in divergence time estimation due to taxon sampling and methodology. Unlike the crown group Papilionoideae origination proposed by Uluer *et al*. (2022), our results suggest a more dynamic evolutionary history of keel flowers with four independent origins that postdate crown group Papilionoideae (Table S11; Fig. S5). Three of these events are nested within the ADA clade (e.g., Dipterygeae and *Myrospermum*), taking place in the Neotropics during the Miocene. The two oldest keel flower lineages originated during the Selandian age of the Paleocene in Papilionoideae and Polygalaceae, though the exact order cannot be confidently established. However, the short interval between these two old events and multiple more recent origins of keel flowers in the ADA and polygalean clades may result from younger lineages inheriting and further consolidating the strong relationship with pollinators already established by older keel lineages (Tucker, 2002; Bello *et al*., 2012; Uluer *et al*., 2022).

In addition to gains, perhaps what is more intriguing is the numerous losses of keel flowers. We identified more than two dozen independent losses in Papilionoideae alone (Table S10). These results suggest that ancestral Fabaceae likely took a generalist pollination strategy, allowing floral symmetry and petal number to be more labile (Arroyo, 1981; Bruneau *et al*., 2014; Sinjushin, 2021). This is exemplified in multiple early branching papilionoids such as *Harleyodendron* R.S. Cowan with rosid-like radial symmetry and lost, reduced, or poorly differentiated petals. These modifications are accompanied by expansion or shifts of pollinators to bats, passerine birds, hummingbirds, and even beetles (e.g., Arroyo, 1981; Pennington *et al*., 2000; Lewis *et al*., 2003). Such pronounced heterogeneity in floral architecture and pollination syndrome mirrors the early evolution of angiosperms and many clades in Gesneriaceae and Ranunculales (Damerval & Nadot, 2007; Martén-Rodríguez *et al*., 2009; Endress, 2010). While up to 28 losses of keel flowers occurred in early branching papilionoids, only three were found within the NPAAA clade, which accounts for more than 70% of species diversity in Fabaceae. All three losses were restricted to the genus *Erythrina* L. (Millettioid-Phaseoloid clade), which is a pantropical genus of ca. 120 bird pollinated, mainly arborescent species (Bruneau, 1996, 1997). Species in *Erythrina* still retain the bilateral floral architecture but differentiate from a typical keel flower in that the stamens and style are exposed rather than protected. Their exceptionally long upper banner petal forms a pseudotubular structure enclosing the other petals, whereas the banner in a typical keel flower folds convexly upward into a boat-shape (Nesom, 2015).

The majority of the NPAAA clade, on the other hand, exhibited significant conservatism in floral form. We hypothesize this to be a joint outcome of ecological pressures imposed by pollinators and herbivores as well as developmental canalization (Bruneau *et al*., 2014; Carvalho *et al*., 2023; Ojeda *et al*., 2019). Although most Fabaceae are bee-pollinated regardless of floral architecture, more reliable and specialized bee associations are developed with true keel flowers in Papilionoideae (Armbruster, 2014; Arroyo, 1981; Fenster *et al*., 2004). The NPAAA clade has diversified largely since the Oligocene under global cooling, aridification, and expansion of grasslands (Estep *et al*., 2014; Herbert *et al*., 2016). The IRLC, Indigofereae, and Robinioid clades are thus strongly associated with herbaceous growth habit in arid to semi-arid temperate regions (Wojciechowski *et al*., 2004; Schrire *et al*., 2005, 2009). Thus adaptation to these habitats may cast additional geographical, environmental, and developmental constraints to limit the breadth of viable pollinators in the NPAAA clade. Moreover, empirical studies from various plant families have also shown that highly developed corollas may protect young buds and fruits from herbivory (Carvalho *et al*., 2023; Herrera, 2010; Gao *et al*., 2019). Finally, developmental canalization may also contribute to homeostasis in floral architecture (Mabberley & Hay, 1994; Leite *et al*., 2015; Ojeda *et al*., 2019). The modularized genetic programs for floral development can produce consistent phenotypes buffered against internal and external influences (i.e., canalization; Moyroud & Glover, 2017). Many ontogenetic modules for keel flowers are shared between Polygalaceae and Fabaceae (Westerkamp & Weber, 1999; Prenner, 2004; Bello *et al*., 2010) and in many cases demonstrate increased canalization among recently diverged angiosperms lineages (Endress, 2011). The conservation of keel flower developmental pathways, including the MADS-BOX and CYC genes for floral symmetry and whorl identity (Singh *et al*., 2013; Zhang *et al.,* 2010), may further contribute to the prevalence of keel flowers in derived Papilionoideae.

### A synergistic multidriver hypothesis for the diversification of Papilionoideae

Evolutionary biologists have been constantly searching for key innovations or environmental stimulants of angiosperm diversity (see reviews from Helmstetter *et al*., 2023; Sauquet & Magallón, 2018). In particular, the direct involvement of floral traits in sexual reproduction has prompted great interest in exploring its phenotypic diversity and link with species diversification (Kay *et al*., 2006). Zygomorphy (Sargent, 2004; Kay & Sargent, 2009), animal pollination (Muchhala, 2006), and nectar spurs (Hodges and Arnold, 1995) have all been suggested to enhance speciation rates. However, a growing body of empirical studies suggested that angiosperm diversification cannot be tied to a single key innovation or major global event (Augusto *et al*., 2014; Vamosi *et al*., 2018). Instead, various combinations of traits, environment, and ecology may have acted to stimulate diversification in different groups (Augusto *et al.,* 2014; Magallón & Castillo, 2009). Such synergistic interaction between multiple, perhaps sequential, intrinsic and extrinsic contributors of diversification rate has been coined as ‘synnovation’ by Donoghue and Sanderson (2015).

In papilionoid legumes, the contrast in species numbers between clades with and without keel flowers motivated our investigation of this phenomenon using comparative methods. The best-supported MiSSE model from our results was trait-free and thus indicated the limited influence of floral architecture on diversification at a global scale. This lack of support for SSE models is likely a conservative conclusion given our taxon sampling (ca. 15% across Fabales) because larger, older, and less well-sampled phylogenies tended to support trait-dependent models (Helmstetter *et al*., 2023). The orders-of-magnitude difference in net diversification rate estimated by the HiSSE model was driven by a handful keel-flowered lineages including the well-characterized radiations in *Lupinus* and *Astragalus* (Fig. 2 a, d). Yet even in these two genera, additional extrinsic and intrinsic factors such as perennial life history and invasion of montane ecosystems have been demonstrated to promote speciation (Drummond *et al*., 2012; Azani *et al*., 2019). Similarly, a lack of correlation between nodulation ability and net diversification rate was reported by Afkhami *et al*. (2018) despite that nodulating legume genera contain three times more species than non-nodulating ones. These results suggest that individual traits such as keel flowers and nodulation are not the sole driver of diversification in papilionoids, but rather constitute a broader confluence of factors that have additively contributed to lineage diversification (Donoghue & Sanderson, 2015).

Indeed, the massive diversity of Fabaceae has been hypothesized to be an outcome of a suite of intrinsic traits including nitrogen fixation symbiotically with soil bacteria (Sprent, 2001; Werner *et al*., 2015; Doyle, 2016; Faria *et al*., 2022), direct chemical defense with alkaloids and canavanine (Bell, 1981; Bisby *et al*., 1994; Kursar *et al*., 2009; Wink, 2013), indirect defense via ant domatia and extrafloral nectaries (Marazzi & Sanderson, 2010; Chomicki & Renner, 2015; Marazzi *et al*., 2019), and likely variation in fruit morphology and seed dispersal mechanisms that have yet to be characterized (López *et al*., 2000; Herrera *et al*., 2019). Besides, recurrent biome shifts and rapid colonization of novel ecological niches also played a significant role in promoting diversification in Fabaceae. Examples include the rapid radiation of the primarily herbaceous IRLC clade in temperate regions (Wojciechowski *et al*., 2004), the dominance of mimosoids in seasonally dry forests (Pennington *et al*., 2009; Ringelberg *et al*., 2023), and the remarkable diversification of Indigofereae in tropical fire-prone savannas (Schrire *et al.,* 2009). Finally, multiple whole genome duplications taking place during early evolution of Fabaceae have also been proposed to buffer mass extinction events and promote genetic and species diversity (Koenen *et al*., 2021; Zhao *et al.,* 2021). Some of these events left a clear signal in our BAMM analysis (e.g., rate shift in mimosoids; node A Fig. 2a) that are uncorrelated with keel morphology. In genera like *Senna, Lupinus*, and *Astragalus*, intrinsic (e.g., extrafloral nectaries, perennial habit) and extrinsic factors (e.g., colonizing Andes) are closely linked with each other to promote diversification (Azani *et al*., 2019; Marazzi and Sanderson, 2010; Drummond *et al*., 2012). Studies of Podalyrieae in the Cape region also concluded that the interaction between fire-survival strategy and soil type preference simultaneously contributed to its diversity (Schnitzler *et al*., 2011). All of this evidence points to the synnovation hypothesis as a more plausible explanation for radiations in Fabaceae.

In summary, while floral architecture may have played a role in the diversification of papilionoid legumes, a combination of intrinsic factors and extrinsic factors were synergistically involved to drive the remarkable diversification of Papilionoideae. This may begin with the accumulation of genetic variation from the paleo genome duplication that occurred along the stem lineage leading to papilionoids (Koenen *et al*., 2021), combined with the origin of nodulation within the clade (Doyle, 2016) and the shifts in floral architecture to bilateral symmetry. Keel flowers may facilitate speciation by enhancing the persistence of species, which might in turn increased the likelihood of experiencing subsequent key innovation or range shifts that promote speciation. Alternatively, keel flowers may represent a phylogenetic inertia for species-rich clades rather than being the primary or casual factor per se of that drove diversification (Blomberg & Garland, 2002).

## CONCLUSIONS

Drawing upon insights from phylogenomics, divergence time estimation, and comparative methods, we present novel insights into the early divergence history and floral evolution of papilionoid legumes. With 1,456 low-copy nuclear loci, we confidently resolved the recalcitrant relationships in Papilionoideae and inferred a Paleocene origin of keel flowers at 59 Ma, predating the earliest fossil record by 3–4 million years. We identified at least six independent origins and thirty-two losses of keel flowers across Fabales, with losses predominantly taking place in the early diversification history of the papilionoids. Ancestral Fabaceae thus likely displayed a more generalist pollination strategy, allowing for greater floral flexibility over time. Although these independent gains and losses coincide with the Miocene diversification of Papilionoideae, our results do not support floral architecture as the sole driver of diversification in Fabaceae. Instead, a confluence of intrinsic and extrinsic factors including nodulation, chemical defenses, and recurrent dispersals into new biomes such as subtropical and temperate regions, likely contributed synergistically to the diversification of papilionoids.

## ACKNOWLEDGEMENTS

This work was supported by grants from the National Science Foundation to MFW (DEB-1853010) and RKJ, TAR (DEB-1853024), the Texas Ecological Laboratory Program (EcoLab) to RKJ, TAR, I-SC, and LC, the Sidney F. and Doris Blake Professorship in Systematic Botany to RKJ, the Stengl Wyer Fellowship to LC, the CNPq (Research Productivity Fellowship no. 314187/2021-9), and FAPERJ (Programa Jovem Cientista do Nosso Estado - 2022, grant no. E-26/200.153/2023) to DC, the National Research Foundation of Korea (NRF) grant funded by the Korea government (MSIT) (RS-2024-00341022) to I-SC, and funds from the College of Liberal Arts and Sciences at Arizona State University to MFW. We thank the TEX-LL, HUEFS, and RB herbaria for voucher deposition, the Desert Legume Program at the University of Arizona for seeds; Janet Davis for kindly allowing us to use her photo of pollinating *Baptisia australis* in Fig. 2; Alessandra Schnadelbach, Henrique Batalha Filho, and Paula Ristow for permitting us to use their labs at UFBA; and George Yatskievych (TEX/LL) for arranging a formal Material Transfer Agreement (Decree number 8772) under the SisGen Cadastro RDC6BE9, which facilitated research activities between the institutions. Comments from the Handling Editor Susana Magallón, three anonymous reviewers, and Michael J. Sanderson have greatly improved the initial manuscript.

## SUPPLEMENTARY INFORMATION

### Supplementary figures

**Figure S1** Histogram of the number of species per loci. The vertical line represents the 30% missing data threshold to select the 1,456 loci in this study.

**Figure S2** Topological congruences and conflicts of Papilionoideae phylogeny inferred from IQTREE (left) and ASTRAL (right). Tangled grey lines indicate conflicting relationships. Branch supports represent nonparametric bootstrap replicates values from IQ-TREE and local posterior probability from ASTRAL, respectively. Only branches that are not maximally supported are labeled.

**Figure S3** Quantification of phylogenetic signal from individual genes. The two alternative placements of the focal clades are illustrated with red or blue, with the red topology being favored by the concatenation analysis. Genes are ordered in descending order based on their support for the red topology. Differences in the log-likelihood scores (ΔlogL) between these two topologies are calculated for individual genes (left) or the concatenated matrix (right). Panels A-E represent five contentious relationships within Papilionoideae defined in Table S3.

**Figure S4** Bayesian inference of divergence time in Fabaceae based on 30 top clock-like loci. Divergence time is shown in millions of years ago (Ma) and the blue bars represent 95% credibility intervals. Eleven fossil calibration points including a constraint for the root age are marked by numbered stars. See Table S5 for a full list of fossils and references.

**Figure S5** Ancestral character reconstruction of keel flowers in Fabales inferred under the marginal reconstruction algorithm in HiSSE. An ultrametric tree of 3,326 Fabales species is inferred from *matK* sequences with branch lengths drawn in proportion to divergence time. Solid and hollow branches indicate the presence and absence of keel flowers, respectively. Events of keel flower gains and losses are marked by orange and magenta circles on the branch, respectively. See Table S11 for age estimations of these events.

### Supplementary Tables

**Table S1** Voucher information and gene assembly information of 79 newly sequenced species in this study. Species name, lineage affiliation, assembly quality, SRA accession number, and voucher information including collection locality are provided.

**Table S2** Species name, lineage affiliation, and number of sampled genes from 208 species from Zhao *et al*. (2021).

**Table S3** Testing alternative phylogenetic placement of Papilionoideae clades under the maximum likelihood framework for concatenated DNA sequences.

**Table S4** List of top 30 clock-like genes with quantified root-to-tip branch length variation, total tree length, and topological similarity to the species tree measured by bipartition score.

**Table S5** Fossil calibration points for divergence time estimation.

**Table S6** Time tree, GenBank ID, and keel flower trait status for the 3,326 Fabales species from the *matK* dataset. Numbers on the phylogeny are divergence time estimation for each node based on treePL.

**Table S7** Secondary calibration points for the treePL analysis. All node ages in million years ago (Ma) are derived from the BEAST time tree and are fixed in treePL. Nodes are defined by the most recent common ancestor of two species indicated in the spreadsheet.

**Table S8** Genus level sampling ratio for Fabaceae. The number of species for each genus was obtained using the powoGenera function from the R package expowo based on the Plants of the World Online database (https://powo.science.kew.org). The species sampling percentage was summarized for each genus and presented along with the URL address from which the POWO data was derived.

**Table S9** Taxon sampling and model parameters for the 11 diversification rate estimation analyses. We used two sets of time-calibrated phylogenies that include either all Fabales or Papilionoideae species only. The BiSSE, HiSSE, CID-2, CID-4, and MiSSE models variously allow for the rate parameters between keel/non-keel states and hidden states to differ. BiSSE, HiSSE, CID-2, and CID-4 models further varied in the number of hidden states between zero and four. The optimum number of MiSSE was determined using the MiSSEGreedy() function based on AIC scores.

**Table S10** Nuclear genome sequencing coverage of 32 whole genome sequencing datasets estimated based on 21-mer frequency distribution. See Table S1 for the species name of each ID.

**Table S11** Summary of independent origins and losses of keel flower in Fabales. Binary ancestral state reconstruction is inferred under the marginal reconstruction algorithm in HiSSE. The timing of these events is determined by the crown group age of involved clades. The precise age of events involving a single species cannot be determined using available data. Therefore, the age of these events is represented by a range from present to their stem group age.

### Supplementary Data

**Data S1** DNA alignments and maximum likelihood phylogenies inferred for the 1,456 low-copy nuclear genes used in this study. The alignments are inferred using the MAFFT-linsi algorithm. The gene trees are reconstructed using IQTREE and branch support is evaluated with 1000 ultrafast bootstrap replication. Species names and their corresponding ID are provided in sp_ID.csv. The list of 672 loci used to generate the concatenated matrix for species tree estimation is provided in G672_len900_sp227.list. Data is available at the Figshare Digital Repository (DOI: 10.6084/m9.figshare.27135984).

**Data S2** Distribution of 21-mer frequency in 32 species with low-pass genome sequencing data. The 21-mer frequency is inferred from Jellyfish v2.3.0 to estimate sequencing coverage.

**Data S3** Gene and site concordance factors of the Papilionoideae phylogeny. The species tree and concatenated alignment of the 672-gene dataset were used as inputs in IQ-TREE to calculate these concordance factors.

## COMPETING INTERESTS

None declared.

## AUTHOR CONTRIBUTIONS

Conception and experimental design, LC, DC, I-SC, RKJ, TAR, MFW. Acquisition of funds, RKJ, TAR, MFW. Field collections, I-SC, DC, HCL, LPQ, RKJ, TAR, MFW. Nucleic acid isolation, I-SC, CL, TAR. RNA isolations, TAR. Transcriptome and genome assembly, CL, LT, BS, LC. DNA alignments and phylogenetic analyses, LC, I-SC, DC, MFW. Data interpretation, LC, I-SC, DC, MFW. Production of figures and tables, LC, DC. Writing the initial draft of the manuscript, LC, DC, MFW, TAR, RKJ. All authors revised and approved the final version of the manuscript.

## DATA AVAILABILITY

The raw sequence data generated and analyzed during this study are available in the NCBI GenBank repository under Bioproject PRJNA1120003. The alignment data supporting the findings of this study are available as supplementary data and have been deposited in the Figshare Digital Repository (DOI: 10.6084/m9.figshare.27135984). The multi-locus phylogeny of Papilionoideae and the dated *matK* phylogeny of Fabales are available on the Open Tree of Life (https://tree.opentreeoflife.org/curator/study/view/ot_2372). The code used for data processing and analysis is openly available on GitHub at https://github.com/lmcai/Papilionoideae_phylogenomics.

## Notes

https://github.com/lmcai/Papilionoideae_phylogenomics

### Competing Interest Statement

The authors have declared no competing interest.

### Summary of Updates

New SSE analyses, major text revision in Introduction and Discussion.

