## Supplementary figures and images for "Well-resolved phylogeny supports repeated evolution of keel flowers as a synergistic contributor to papilionoid legume diversification"

### Fig. S1

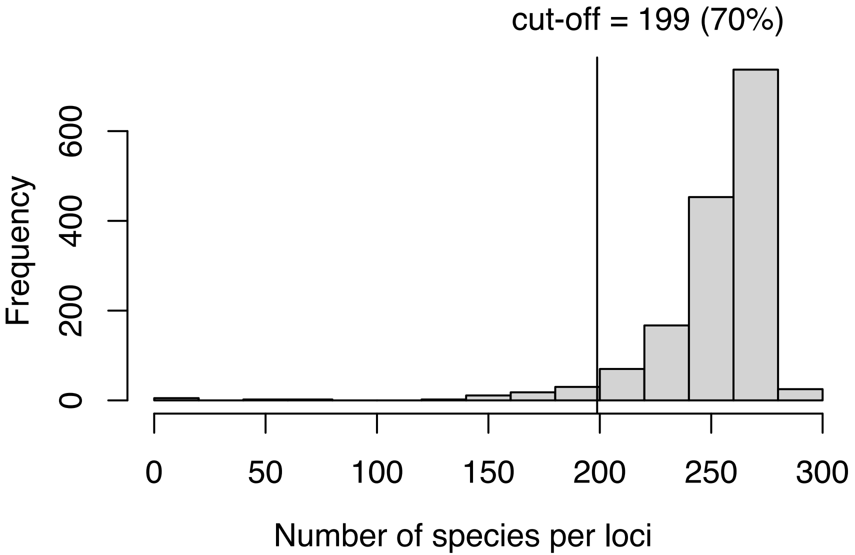

### Fig. S2

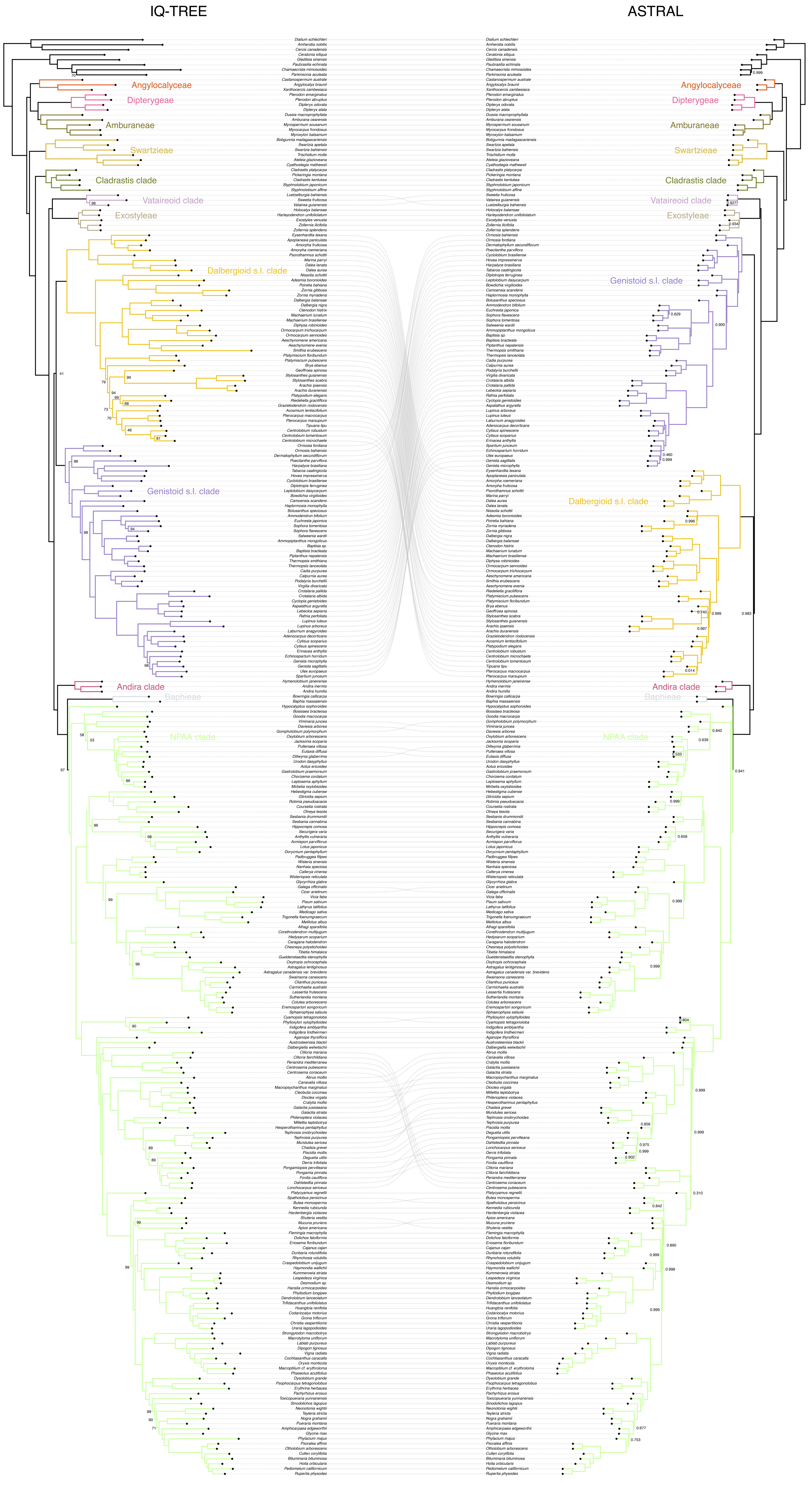

### Fig. S3

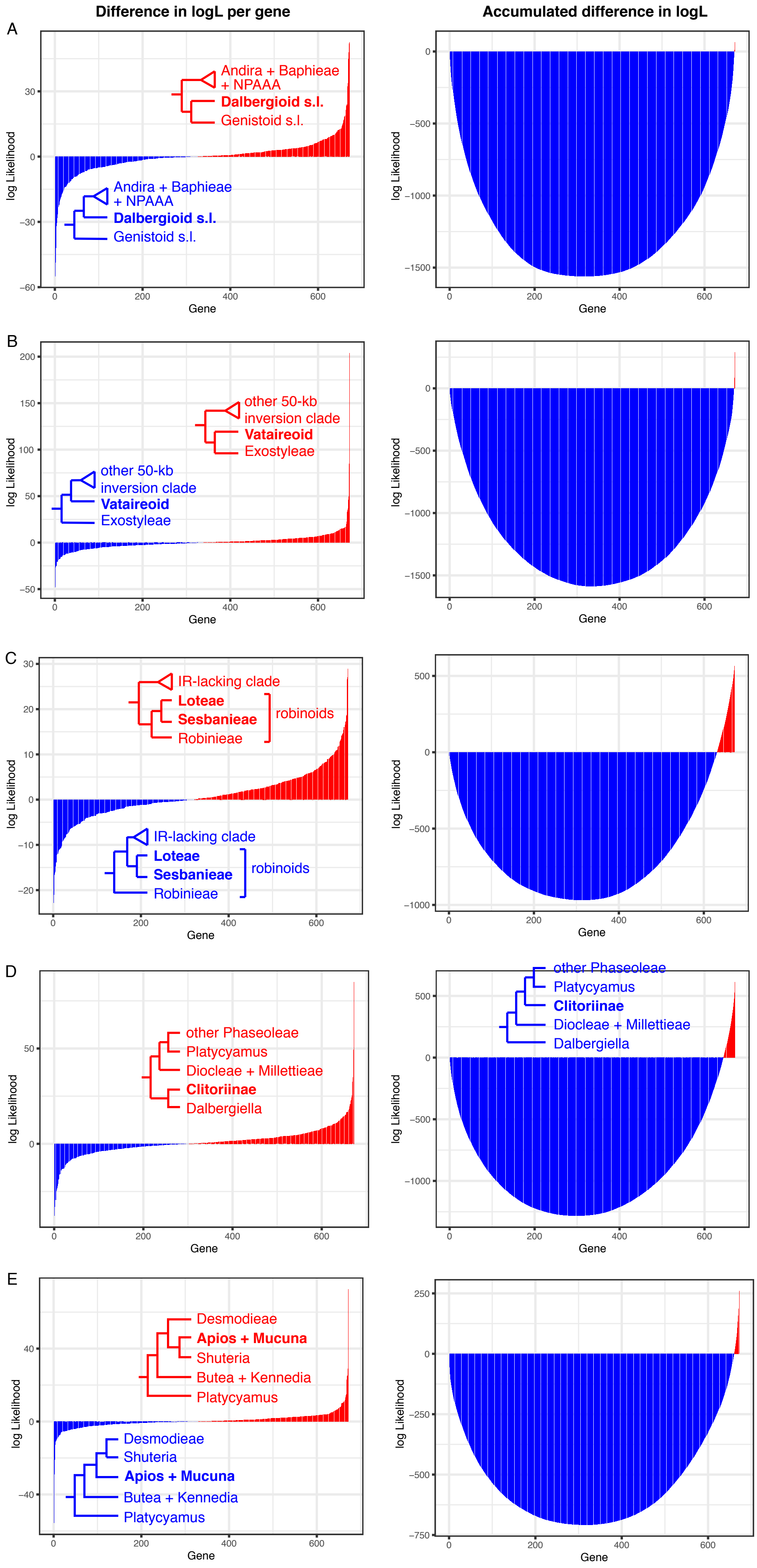

### Fig. S4

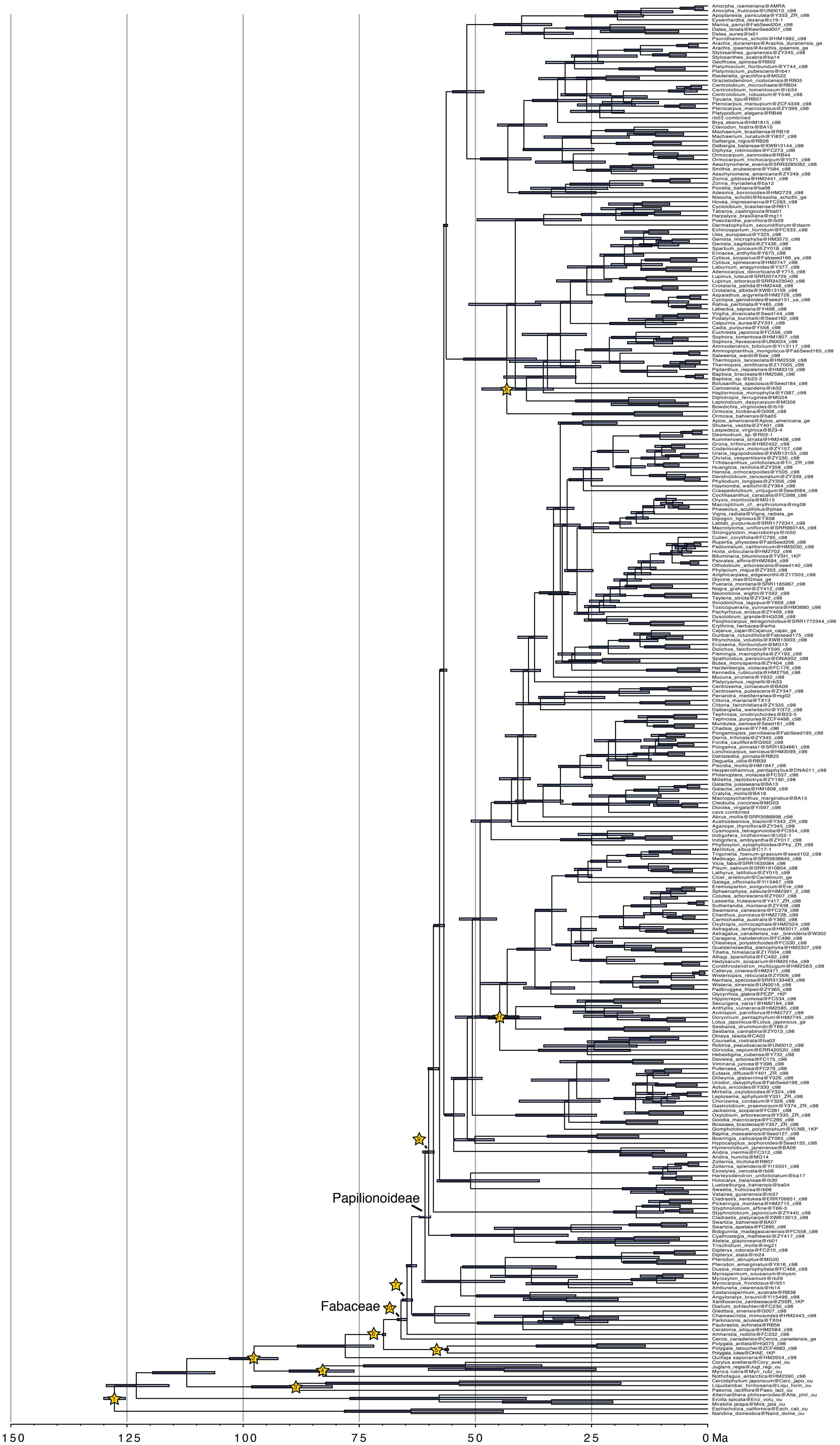

### Fig. S5

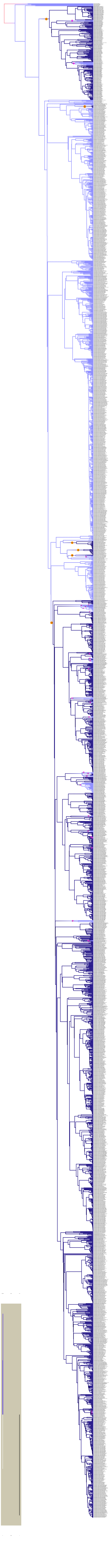
